# An Adaptive Geometric Search Algorithm for Macromolecular Scaffold Selection

**DOI:** 10.1101/099762

**Authors:** Tian Jiang, P. Douglas Renfrew, Kevin Drew, Noah Youngs, Glenn Butterfoss, Dennis Shasha, Richard Bonneau

## Abstract

A wide variety of protein and peptidomimetic design tasks require matching functional three-dimensional motifs to potential oligomeric scaffolds. Enzyme design, for example, aims to graft active-site patterns typically consisting of 3 to 15 residues onto new protein surfaces. Identifying suitable proteins capable of scaffolding such active-site engraftment requires costly searches to identify protein folds that can provide the correct positioning of side chains to host the desired active site. Other examples of biodesign tasks that require simpler fast exact geometric searches of potential side chain positioning include mimicking binding hotspots, design of metal binding clusters and the design of modular hydrogen binding networks for specificity. In these applications the speed and scaling of geometric search limits downstream design to small patterns. Here we present an adaptive algorithm to searching for side chain take-off angles compatible with an arbitrarily specified functional pattern that enjoys substantive performance improvements over previous methods. We demonstrate this method in both genetically encoded (protein) and synthetic (peptidomimetic) design scenarios. Examples of using this method with the Rosetta framework for protein design are provided but our implementation is compatible with multiple protein design frameworks and is freely available as a set of python scripts (https://github.com/JiangTian/adaptive-geometric-search-for-protein-design).

## 1 Introduction

The field of protein design has advanced tremendously in the recent decade in scale, accuracy and the number of types of design tasks carried out by practitioners. Early successes in protein design focused on protein fold design (including novel folds)[1] and hyper-stabilisation of proteins[2]. The redesign of protein-protein[3] and protein-DNA[4] interfaces allows for functional rewiring of key biological networks. More recently, protein engineers have turned towards the redesign of protein active sites and smaller functional patterns that demand sub-angstrom accuracy in the positioning of key side chains. Recent works include both the engraftment of known active sites onto new scaffolds[5] as well as the engraftment of novel active sites (derived from quantum mechanical modeling of desired reactions and corresponding transition states)[6] onto new scaffold proteins. In these enzyme design applications, active site patterns can become quite large as substrate binding, reaction mechanism, and surrounding environment are considered. Methods for matching known and predicted functional sites onto large libraries of potential scaffolds (proteins, nucleic acids and synthetic peptidomimetics for example) are needed to enable enzyme design (and other related design tasks involving functional site or hotspot transplantation).

The earliest geometric matching applications in bioinformatics were aimed at matching whole sub-structures to find substructures that indicated a likelihood of shared function or distant homology[7]. In many cases these algorithms searched for contiguous regions and were the structural analog of sequence alignment algorithms (both gapped and ungapped). Early uses included protein function prediction, analysis of structure prediction and evaluation of new algorithms[8–10]. Early works also included innovative uses of geometric hashing to extract 3D functional motifs from protein structures[11]. Here we focus on the use of geometric search for the purpose of biodesign, rather than prospecting or annotation.

Geometric searches in biodesign and bioinformatics contexts having similar motivation to the work described here have used combinations of geometric hashing, side chain conformation libraries and other heuristics that have typically limited the number of elements in any given pattern. Fleishman et al. computationally designed a protein to bind hemaglutinin (HA) targeting a conserved region on the stem[12]. They first identified residues that were likely to be hotspot residues by docking the single amino acid onto the HA stem region and calculating a binding energy. Next, they built inverse rotamer libraries for residues with good binding energies, then used the residues as anchor sites on which to dock protein scaffolds. The protein scaffolds were selected from known proteins not known to bind HA and were filtered for high shape complementarity with the HA target region. A low resolution docking procedure was used to simultaneously optimize the HA-scaffold binding energy as well as the scaffold’s ability to accommodate anchor residues. Scaffolds that showed good binding energies with the satisfied hotspot residues were used as the starting point for a second round of docking and designing to optimize residues outside of the hotspot residues.

In addition to these examples of geometric search-driven enzyme design, there are several examples in the field of biomimicry with synthetic oligomeric foldamers and short peptidomimetic scaffolds. Here the objectives vary considerably: in active-site mimicry, interface binding, metal binding and surface adhesion[13] are a few of the diverse pepdiomimetic design tasks. The set of oligomeric scaffolds that have protein-like side chain take-off angles is quite diverse; examples include linear peptoids[14], oligooxipiperizines (OOPs)[15], HBS helices[16], cyclic peptides[17] and peptoids[18], β-peptides[19] and hybrids thereof. A key application here is the mimicry of interfacial hotspots where a small number of side chains scafolded by a single secondary structure element comprise a significant fraction of the binding energy[20]. In these cases moving these groups of side chains to a new, non-protein, scaffold with synthetically restricted backbone degrees of freedom and reduced atomic mass is a viable route to inhibiting protein interactions. Drew et al. show that by grafting 4 side chains from a restricted segment of sequence onto a four subunit OOP scaffold resulted in low nanomolar inhibitors of two key protein-protein interactions (p53-MDM2 and p300-Hif1α)[21]. The first step in this work was using a geometric search to dock the OOP scaffold into the binding site such that side chain takeoff angles were compatible with those observed for three hotspot residues observed in the structure (predicted to comprise the majority of the binding energy). After this geometric search was used to instantiate a starting pose, the Rosetta design procedure (with modifications for both NCAA side chains and the OOP backbone) was used to optimise binding and inhibition of the endogenous protein-protein interaction, resulting in low nanomolar inhibitors of both complexes. In both of these cases, the geometric match steps were based on inverse rotamers and were prohibitively expensive, limiting the search to only small peptidomimetics.

Drew and Renfrew et al. previously demonstrated the incorporation of several non-peptidic backbone chemistries in the the macromolecular modeling suite, Rosetta[22]. There are many additional abiotic foldamer and peptidomimetic backbone bones[23] that are amenable to such treatment. Determining which foldamer backbone (or hybrid) chemistry) is the most compatible with a given interface will become a bottleneck as the number of synthetically accessible scaffolds for biomimicry continue to increase.

Here we describe a new method combining octrees (a data structure that maps regions of 3-dimensional space to nodes in a tree) and a novel adaptive search that results in a significant performance gain for the applications described above. Key innovations include the ability to weight interaction/pattern components by energy and the adaptive nature of the search, which both increase efficiency and allow for specification of allowable error (per component of the template pattern) and number of mismatches. We pose the problem by describing a typical problem setup. We then describe our core algorithm. Lastly, we describe applications to protein and peptidomimetic design tasks like those described above.

## 2 Methods

### 2.1 Problem Setup

Here we describe a method that, given a set of side chain functional groups that are fixed in space, will find a molecular *scaffold* among a library of scaffolds that will accommodate those fixed *functional groups*. Here, we use the term functional group to describe the terminal atoms of a side chain, i.e. those atoms whose position will remain fixed relative to one another during the rotation of the *χ* angles of the side chain. These would include the phenyl, imidozol, and guanadinium groups of phenylalanine, histidine and arginine respectively, but also the four terminal carbons of leucine (Cβ, Cγ, Cδ1, Cδ2) and the hydrogens that branch from them. A molecular scaffold is defined generally as any molecule from which designable side groups could branch.

A given scaffold will typically have varying degrees of freedom and these degrees of freedom will therefore define the scaffold’s ability to accommodate fixed functional groups. Practically, different scaffolds will have different degrees of flexibility at different positions and this will drive our definition of allowable error of matching. For a peptide the degrees of freedom are the φ and ψ angles on the backbone and *χ* angles in the side chains. Peptidomimetic scaffolds will have different/additional degrees of freedom. For example, for a peptoid we must also consider the preceding-ω angle which potentially allows for greater diversity of side chain Cα-Cβ bond vectors for a given sequence. Alternatively, an oligooxopiperazine (OOP), which has cyclic constraints between neighboring residues, is theoretically much more restricted in its ability to accommodate fixed functional groups but also has a reduced entropic cost upon binding a target.

Our approach to interface design is be part of a two-step process. In the first step, we consider the most influential energies and conduct an efficient geometric search to eliminate all the impossible designs. In a second step, designs that passed the quick initial screening are further refined using Rosetta, potentially introducing additional mutations. This two-step process efficiently saves all the time that the majority impossible designs would take to be evaluated by Rosetta. The second step in this process

### 2.2 Current Hotspot Matching Algorithm

For comparison we adapted the approach of Flieshmann et al. Our implementation of scaffold matching for proteins is quite similar to the above described approach. This approach is broken into three stages as follows:

A. Identify hotspot residues at the interface of a protein interaction subject to the constraint that the residues are not part of the target protein. Hotspot residues are generally chosen based on high ΔΔG values in their alanine scans. Such residues are often responsible for the protein interaction’s binding affinity.
B. For each hotspot residue, generate an inverse rotamer library which specifies high probability orientations of backbone atoms and other atoms not included in the residue’s fixed function group. The inverse rotamer library defines possible connection points to the molecular scaffold of interest.
C. For every designable residue position on a given molecular scaffold a. identify a primary hotspot residue generally chosen as the residue with the highest ΔΔG value from the alanine scanning results b. Align the designable residue position on the scaffold with an inverse rotamer in the library of primary hotspot residue inverse rotamers c. Sample the scaffold’s degrees of freedom to minimize an energy function as well as the distance between remaining designable residue positions on the scaffold and the remaining hotspot residues. In practice, a distance constraint is placed between the atoms at the designable residue positions on the scaffold and the corresponding atoms in the inverse rotamers of hotspot residues and is incorporated into the energy function to evaluate the entire system. d. Save lowest energy conformations and filter for scaffolds that accommodate multiple hotspot residues.

### 2.3 Overview of Adaptive Geometric Search Algorithm

Here we employ octrees as the core data-structure for our algorithm[24]. A cubic volume, with sides of length *l*, centered on a point, *p*, can be subdivided in to eight cubes with sides of length *l*/2, that share *p* as a vertex. Each of these eight cubes can be further subdivided in to eight more cubes each with side of length *l*/4, and so on. This decomposition of 3D space lends it itself toward a tree like representation called an octree. Octrees are tree structures whose nodes correspond to 3D cubes embedded in a hierarchically subdivided overall 3D space and each deepening level of the tree describes an increasingly smaller volume of space. Each node has eight children by subdividing each side of the cube by the middle in the *x*, *y* and *z* dimensions. All the 3D objects are stored in the leaf nodes in the octrees. Octrees have various stopping criteria to stop the tree from splitting including thresholding based on the number of objects in a node, i.e. the octree splits only the nodes containing more than a certain number of objects. For our problem the 3D objects are points in 3D space and the stopping criterion is the minimum cube length *ℓ_s_*. That is, the octree splits a node only if its corresponding cube has sides of length at least 2*ℓ_s_*. Moreover, all empty nodes, i.e. nodes whose corresponding cubes contain no points, in the octrees are discarded.

To search for desirable configurations, the algorithm first samples points from each manifold (corresponding to a take-off point) and then builds an octree for each manifold based on these sample points with the stopping criterion of the minimum cube length *ℓ_s_*. Then the algorithm compares two octrees at a time by searching adaptively in the cubic regions that pass the necessary condition A (see below). We call a pair of cubes that pass the necessary condition A a “possible pair”. The algorithm finds all the possible cube pairs at each level until it ends up with the set of all possible pairs of leaf cubes. Then the sufficient condition A (see below) is tested on all these pairs of leaf cubes to determine whether to accept or reject all the pairs of points inside them. At the end all the pairwise desirable cubes are combined through a matrix product.

### 2.4 Establishing Necessary and Sufficient Conditions for matching

Our overall strategy is to enumerate all possible residue positions (when there is a choice) and amino acid assignments to these residues and then to use the adaptive geometric algorithm to determine whether the resulting rotamers at those positions have the proper geometry. Thus the adaptive geometric algorithm is the “inner loop” of the computation. For this inner loop to be efficient, it must swiftly filter away impossible geometries (theorem 1 below) and identify promising ones (theorem 2 below).

Mathematically, the adaptive geometric algorithm efficiently searches for a certain n-polygon among n sets of points in 3D space given an error tolerance and an approximation margin. This general scheme is required for all the applications introduced above and evaluated in the Results section. Given a target polygon 𝒫 = {*P*_1_, *P*_2_,…, *P_n_*}, a tolerance *∊_T_* ≥ 0 and one edge (*P_i_*, *P_j_*), let 𝒞_*j*_, 𝒞_*j*_ be two nonempty cubes with size *ℓ* and the distance between their centers *d*, where *i*, *j* ∈ [1, 2,…, *n*], *i* ≠ *j*. Then we have the following theorems that help us determine which cubes could possibly match that edge. That is, the theorems provide acceptance and rejection criteria for pairs of cubes from different manifolds (where each manifold corresponds to, for example, a take-off residue from a backbone). The first theorem provides a rejection criterion.

#### Theorem 1.

***I****f* 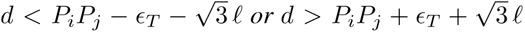, *then there are no pairs of points* (*G*, *H*) ∈ 𝒞_*i*_ × 𝒞_*i*_ *such that* | *GH* − *P_j_P_j_* | ≤ *∊_T_*.

*Proof.* See Appendix A.

Theorem 1 suggests a “necessary condition” for any two cubic regions on the same level of the trees to contain any desirable pairs of points. We are going to call it the “necessary condition 1” in the future to refer to the condition defined in Theorem 1. If two cubes do not satisfy the conditions of this theorem, they are not going to match the edge, and will be rejected. That’s why we consider this to be a rejection condition. By contrast, we have the following “sufficient condition 2” for all pairs of points from two leaf cubes to be desirable (an acceptance condition).

#### Theorem 2.

***I****f* 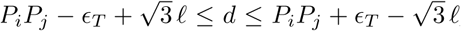, *then all pairs of points* (*G*, *H*) ∈ 𝒞_*i*_ × 𝒞_*i*_ *satisfy* |*GH* − *P_j_P_j_* | ≤ *∊_T_*.

*Proof.* See Appendix A.

Notice that the condition of Theorem 2 can hold only when 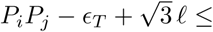 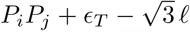 or when 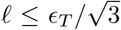. Because the leaf cubes of the octrees must have length *ℓ_T_* ≤ 2*l_s_*, we require 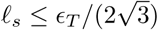.

Let *t_i_* be the octree generated from manifold *𝓐_i_* for *i* = 1, 2,…,*n*. Algorithm 1 gives the pseudo code of the adaptive geometric search algorithm.

#### Algorithm 1 Adaptive Geometric Search ({𝓐_1_, 𝓐_2_,…, 𝓐_*n*_}, 𝓟, *∊_T_*)

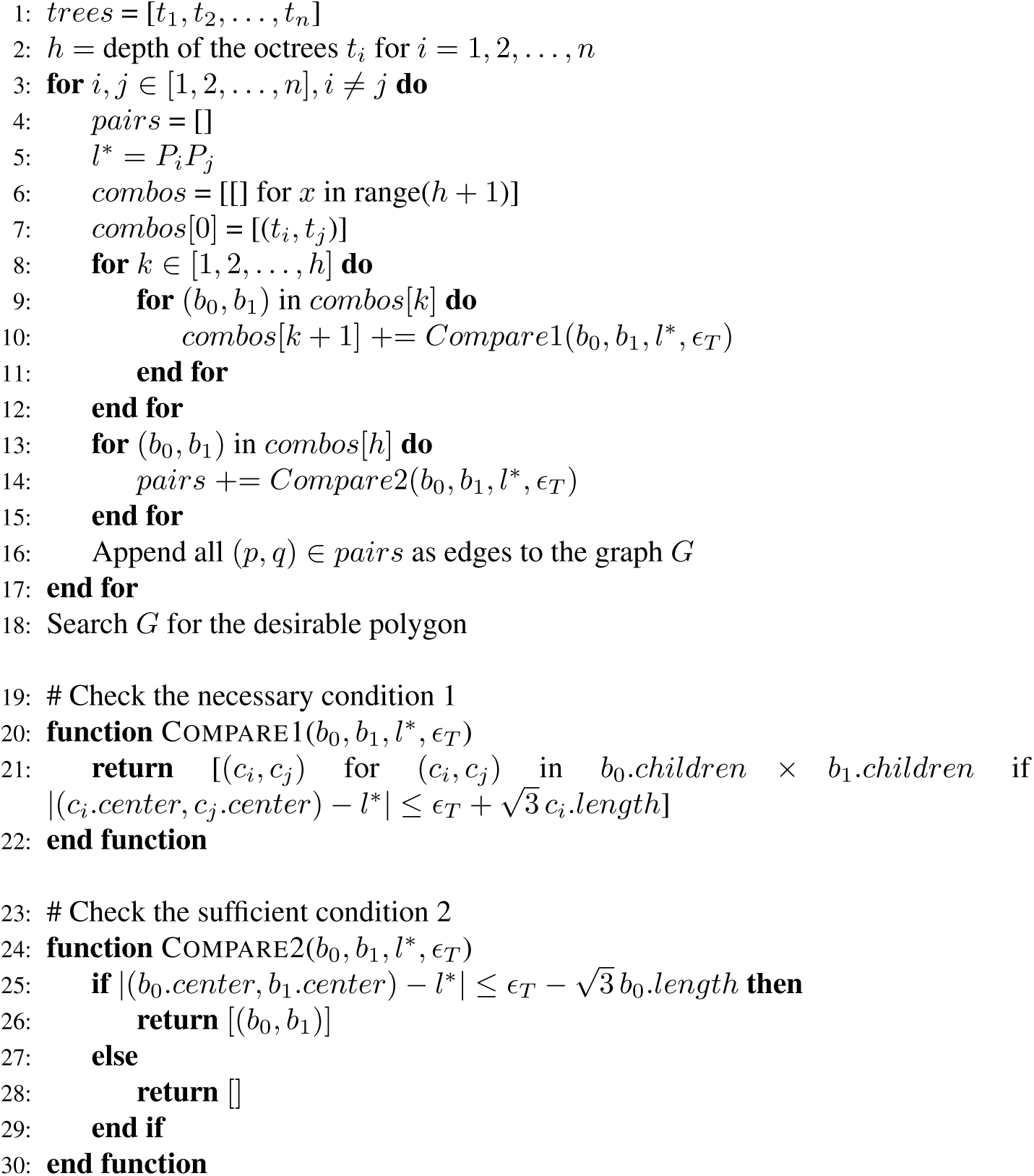

### 2.5 Algorithmic Complexity

The adaptive geometric search algorithm has three parts, building the octrees, adaptively searching every two octrees and the graph search. Let *N* be the number of sample points from each manifold. For convenience we build all octrees with the same initial cube length *ℓ*_0_. The time complexity of building an octree with initial cube length *ℓ*_0_ and minimum cube length *ℓ_s_* is 𝒪(log_2_(*ℓ*_0_/*ℓ_s_*)*N*).

Next we compute the time complexity of the adaptive search between any two octrees (without loss of generality) called *t*_1_, *t*_2_. Let the corresponding polygon edge length be *l*\*.

#### Theorem 3.

***I****f we set* 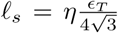 *for any* 0 < *η* < 1, *then the adaptive geometric search algorithm 1 returns all the pairs of points whose distances are within the set* [*ℓ*\* − (1 − *η*)*∊_T_*, *ℓ*\* + (1 − *η*)*∊_T_*], *and some but possibly not all the pairs of points whose distances are within the set* [*ℓ*\* − *∊_T_*, *ℓ*\* − (1 − *η*)*∊_T_*) ∪ (*ℓ*\* + (1 − η) *∊_T_*, *ℓ*\* + *∊_T_*].

*Proof.* See Appendix A.

#### Lemma 4.

*Set* 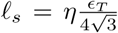 *for some* 0 < *η* < 1. *Then for any cube* 𝒞_1_ *in an octree t*_1_, *there are at most* 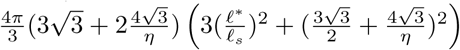 *cubes* 𝒞_2_ *on the same level from another octree t*_2_ *such that* (𝒞_1_, 𝒞_2_) *are possible pairs, that is, they satisfy the necessary condition 1*.

*Proof.* See Appendix A.

#### Theorem 5.

*Recall that ℓ*_0_ *denotes the initial cube length and the minimum cube length* 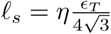. *Let n_m_ be defined as in Lemma A. Then the time complexity of the adaptive search part of Algorithm 1 is* 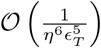.

*Proof.* See Appendix A.

Now we consider the last part of the algorithm, the graph search. Let *s_ij_* be the number of possible leaf cube pairs between octrees *t_i_*, *t_j_* for *i*, *j* ∈ [1, 2,…, *n*], *i* < *j*. We view the leaf cubes as vertices and possible pairs of them as undirected edges in the graph. If we want to produce all the desirable n-tuple cubes, then by induction it’s easy to see that the upper bound on the time complexity is 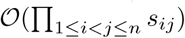.

In practice we can do much better. Consider building a directed graph by giving directions to the edges to form a n-cycle of groups of cubes from *t*_1_, *t*_2_,…,*t_n_*. Finding strongly connected components in this directed graph first would in most cases greatly reduce the search space with only a linear cost 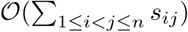.

In summary we state the total time complexity of the algorithm.

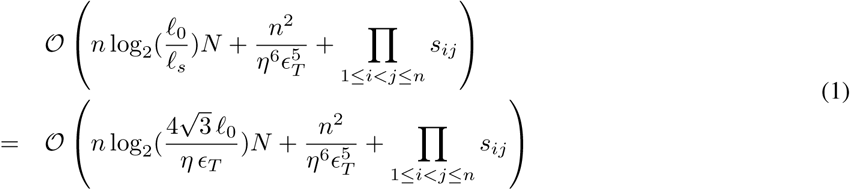

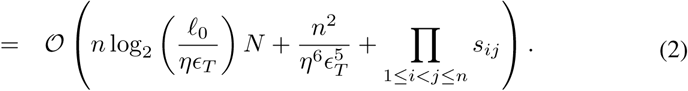

In practice we usually search for a triangle or a 4-sided polygon as the target polygon, i. e. *n* = 3 or 4. When *n* = 3, depending on the parameters *η*, *∊* and *N* the computation time varies but all three terms in the complexity formula (4.1) are typically of the same order. When there are large numbers of possible pairs *s_i_*’s and/or *n* = 4, the term 𝒞(*S*) in the last term of the complexity formula (4.1) becomes the dominating term. The number of results *s_ij_*’s can be further reduced when we take optimal dihedral angles instead of uniform sampling from [0, 2*π*].

## 3 Results

Our algorithm can be applied to many different problems in macromolecular modeling and design. In essence, it efficiently solves the problem of searching for a certain *n*-polygon among *n* sets of points in 3D with error tolerance *∊* and an approximation margin *η*. We present three use cases where our algorithm’s improved efficiency (run times that are in some cases many thousands of times faster than previous approaches) improves the scaling of the overall task, enabling the use of larger template/target structural patters.

### 3.1 Scaffold Matching: designing OOPs to inhibit MDM2-p53 interface

Protein-protein interactions (PPI) mediate many cellular functions and a small number residues that make significant contributions to the the binding affinity of the PPI (deemed “hotspot” residues) in turn underlay these protein interfaces. Design tasks aimed at protein interfaces abound, for example Fleishman et al. previously designed a influenza hemaglutinin binder. Interest in using smaller, easy to synthesize, non-proteolyizable macromolecules (called foldamers) as potential therapeutic candidates continues to rise as these systems continue to become more computationally and synthetically accessible to a broader community. Foldamer backbone chemistries abound and finding a foldamer backbone type that is well matched to a particular set of interface hotspot residues interface will prove to be a future challenge. Here we describe the recapitulation of a OOP foldamer scaffold designed by Drew and coworkers that mimics P53 and can disrupt the P53/MDM2 interaction (Fig. 1D). Three hotspot residues on P53 contribute the majority of the binding affinity for MDM2 (Fig. 1A).

**Fig. 1:**
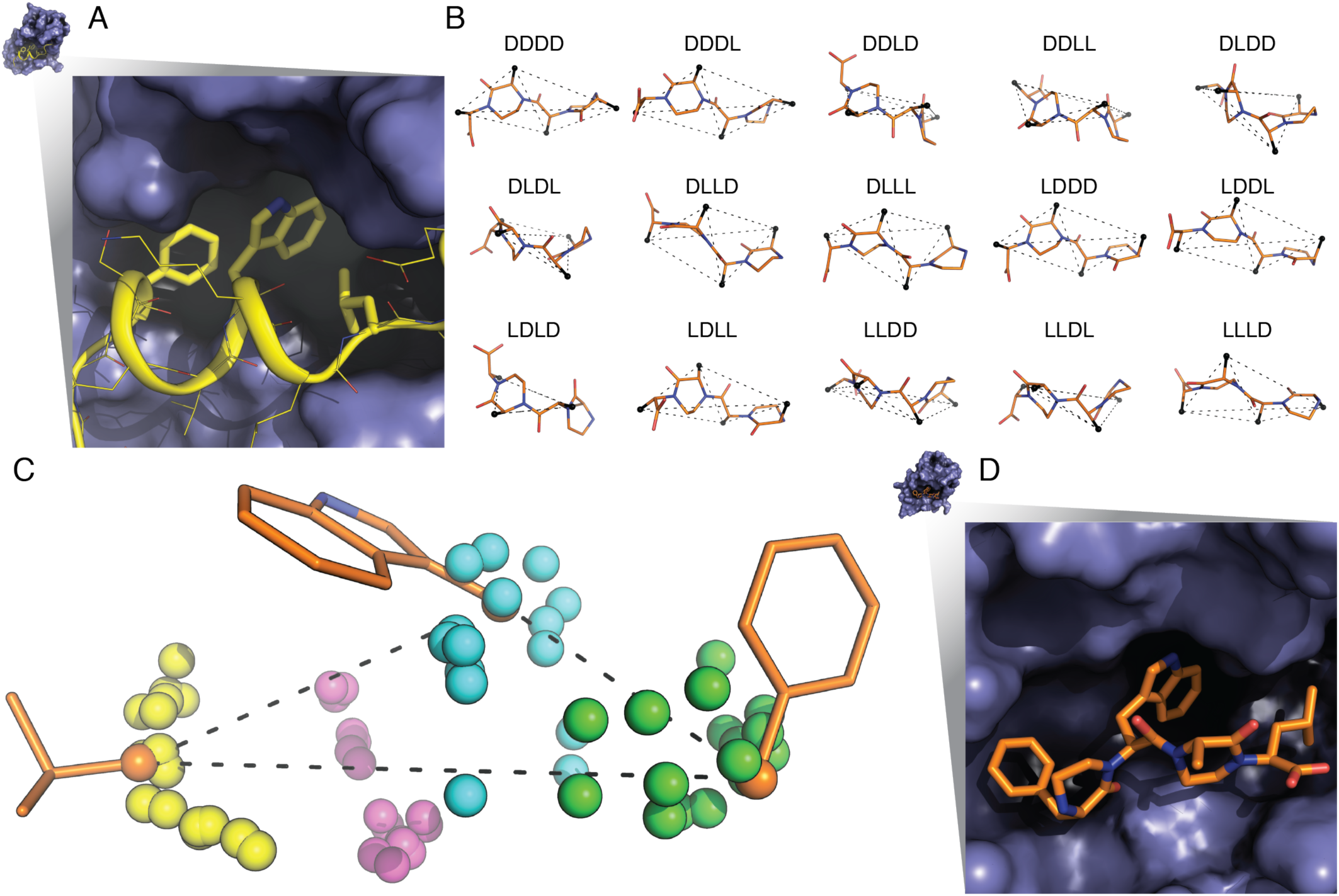
(A) The P53 (yellow) and MDM2 (blue) interface showing phenylalanine, tryptophan, and leucine hotspot residues. (B) Fifteen of the sixteen OOP backbone scaffolds fit to hotspot residue stubs. Scaffolds combinatorially sample the L or D enantiomers of the four residues that comprise the OOP scaffold. Each backbone has four Cβ atoms (black spheres) and thus four possibly matching triangles indicated by dashed lines. (C) The P53 hotspot residue stubs (orange). In this work, each hotspot residue has two χ dihedral angles resulting in a single fixed Cβ (orange spheres) triangle (dashed lines). Hotspot residues with additional χ angles would produce multiple triangles. Colored spheres show potential Cβ atoms from the OOP scaffolds for the first (green), second (cyan), third (magenta), fourth (yellow) residues in the scaffold. (D) The LLLL-OOP scaffold (orange) designed by Drew and coworkers and correctly identified by the algorithm bound to MDM2 (blue).

There are two parts of the algorithm. In step 1, we search through all possible backbones for a matching triangle to the target triangle. In step 2, for every match result from step 1 the connecting atom’s bond angles are checked against the optimal bond angle. If a match passes step 2, it’s returned as a final result. Otherwise we continue the iteration in step 1.

The target triangle is made up of Cβ’s of the hotspot residues (Fig. 1C). The algorithm simply searches through the possible take-off position combinations, four triangles in this example (Fig. 1B), from every backbone for a match in shape within the error bound. Notice that in this case all Cβ’s are fixed due to the short lengths of hotspot residues. With longer hotspot residues, there will be a manifold of all the possible Cβ’s for each hotspot residue. For every possible triplet of take-off positions, there are eight possible D and L entantiomers. So, for each of these 32 possibilities, we apply adaptive geometric search to find all the matches.

Once we have the matching shapes, we calculate the corresponding matrices *R*’s of rotation and translation such that after applying these transformations *R*’s backbones are connected onto the hotspot residues at atoms Cβ’s. Finally we just check if the bondangles at the connecting atoms (eg. N, Cα, and Cβ for leucine) are within some error bound to the optimal bond angles.

#### Algorithm 2 Scaffold Match ({*A*_1_, *A*_2_,…, *A_n_*}, {*P*_1_, *P*_2_,…, *P_m_*}, *δ*, *δ_A_*)

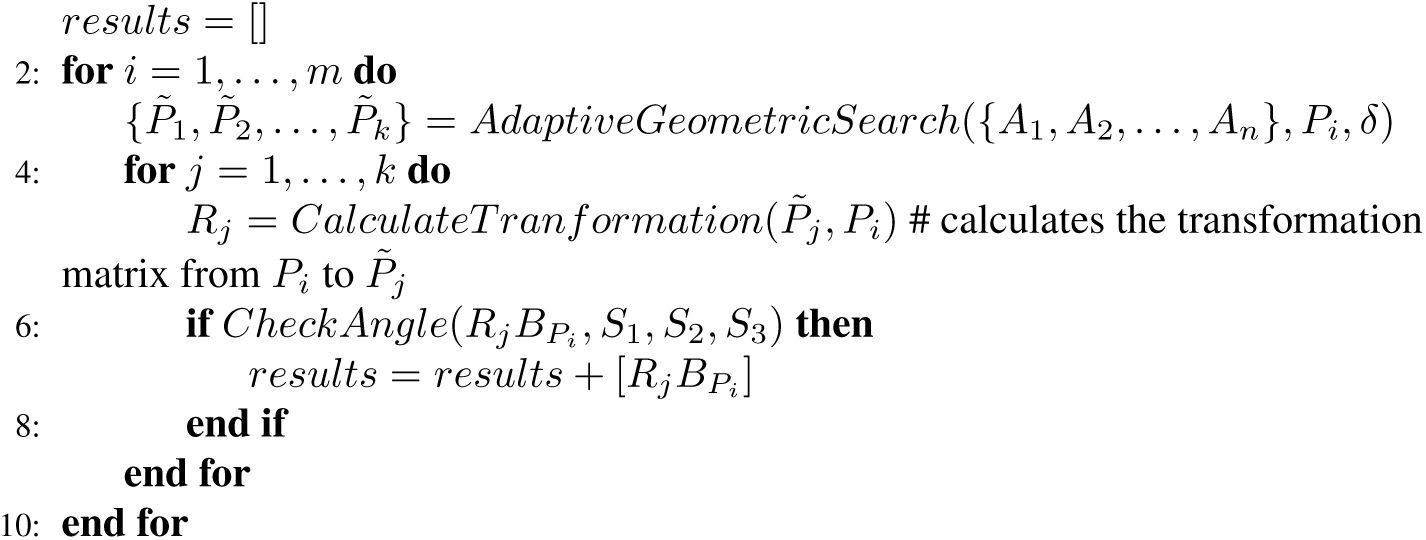

In this example let *A_i_* be the manifold of possible positions of the connecting atom on the *i*-th hotspot residue. For example, in Fig. 4B points in colors are sampled from manifolds *A*_1_, *A*_2_ and *A*_3_ respectively. Let *P_j_* be the *j*-th polygon of the backbone take-off position combination and for example, there are 4 × 17 of them in Fig. 4C. Let *B_P_* denote the atoms positions matrix corresponding to the backbone where the target polygon *P* comes from. Let *S_i_* denote the atoms positions matrix for the *i*-th residue. Let *δ* be the distance error bound and *δ_A_* be the angle error bound. Then we describe in peudocode Algorithm 2. Let 𝒞 denote the time complexity for adaptive geometric search. Recall in Algorithm 2 that *m* is the number of target polygons from backbone take-off site combinations. Then the time complexity of the scaffold matching algorithm is *𝒪*(*𝒞m*).

In the search process we scored all the possible matches by the root mean square deviation (RMSD) values for both shape match and angle match in Fig. 2. Our algorithm picked the candidate at the origin in this plot (this being identical to the correct conformation that led in Drew et al. to low nanomolar inhibitors of this interface). In Fig. 1D we show this best design for the OOP backbone of the hotspot residues. Testing this code as part of an Rosetta OOP-design protocol shows its energy score is a low 4.67 with a potential energy score 4.59 after further minimization, which means this OOP-protein interface is likely very stable (verified experimentally in Drew et al). The run time for the initial geometric search (step on in this design protocol) is 0.02 ~ 0.12 seconds using the algorithm described herein, whereas running the same design and producing the same results using the previously described Rosetta codes (the scripts from Drew et al.) takes ~ 18 minutes (a speedup of greater than 9,000 fold).

**Fig. 2:**
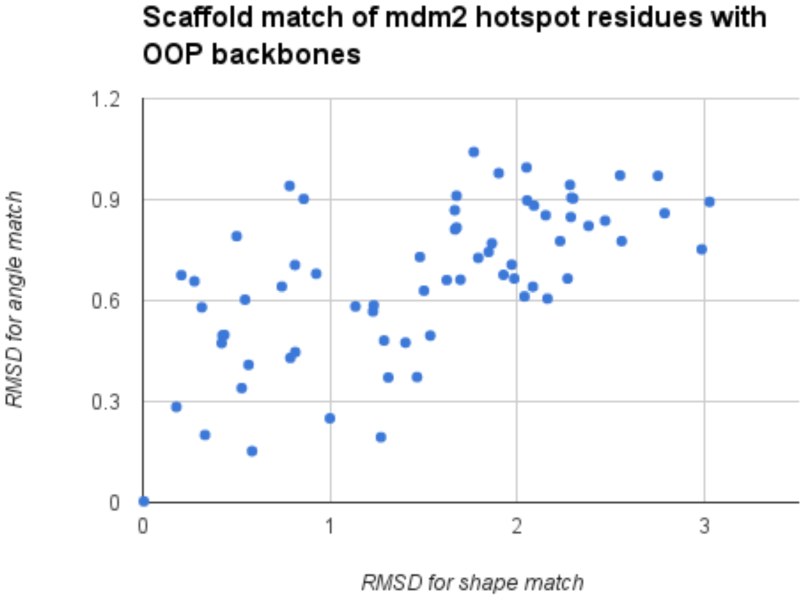
RMSD (root mean square deviation) of all possible OOP backbone matches with the hotspot residues side chain positions. The candidate at the origin is a perfect match for both (shape and angle) to the hotspot residues we aim to minimize (use as a template for design) and is analogous to a template used in previously reported successful experimental designs.

### 3.2 Peptoid design: design of new metal binding sites

Proteins and other macromolecules often coordinate metal ions to aid conformational stability or carry out chemical reactions. Proteins that bind Zn^2+^ ions often use four residues (most often histidine, cystine, or aspartic acid) to coordinate the zinc ion in a tetrahedral arrangement[25]. We next tested our algorithm by designing a peptoid design for capturing zinc ions. The binding sites we target in this example are three sulphur atoms lying on the vertices of an equilateral triangle. A “6-mer” peptoid macrocycle was used as a template backbone (Fig. 3A).

**Fig. 3:**
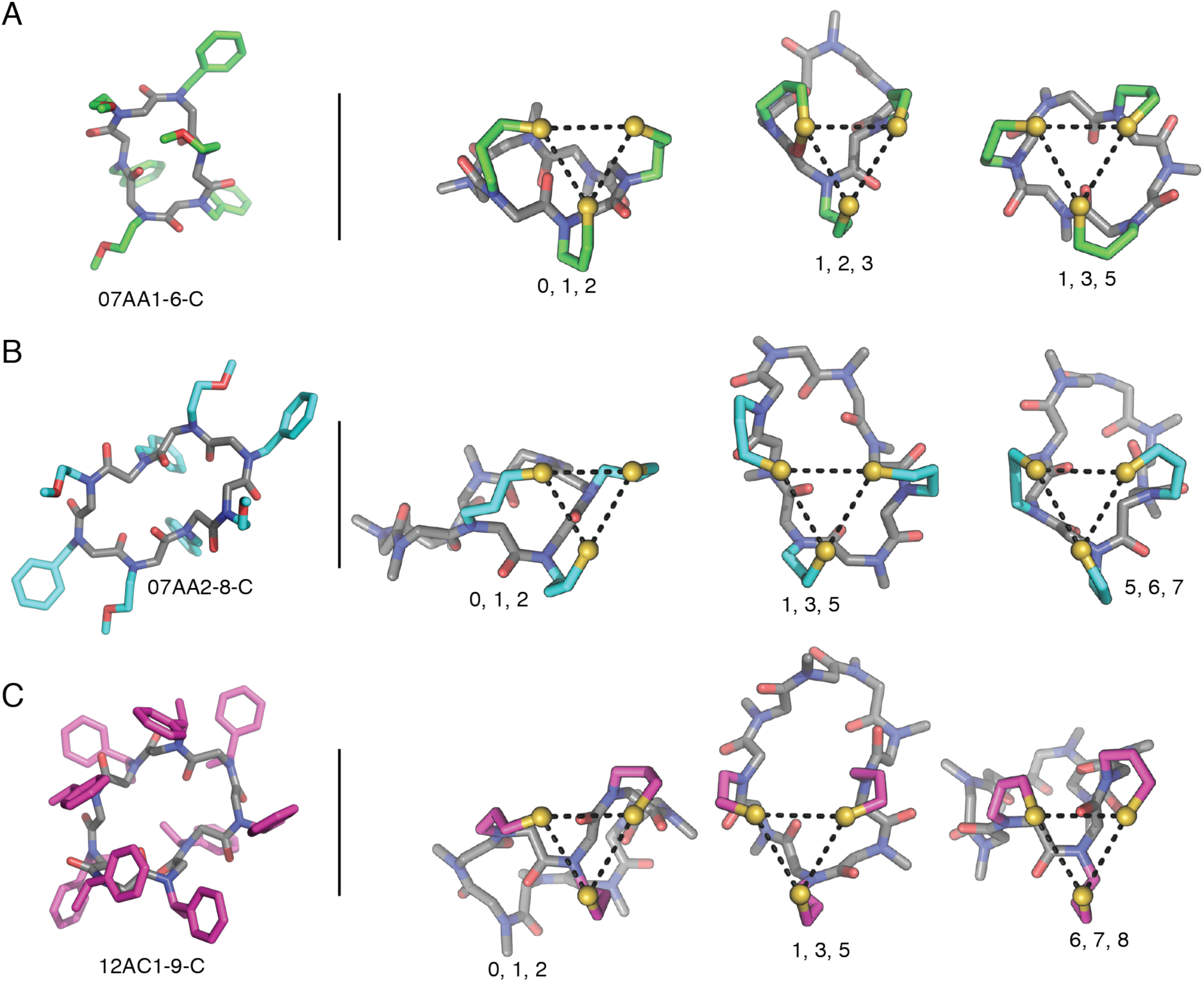
Experimentally determined peptoid macrocycle structures and representative examples of low energy matches for the (A) 07AA1-6-C (B) 07AA2-8-C and (C) 12AC1-9-C peptoid macrocycle backbone scaffolds. Numbers under representative examples indicate residue position of 3-aminopropyl-1-thiol side chain.

A six residue cyclic peptoid composed of alternating sarcosine and 3-aminopropyl-1-thiol side chains. The cyclization directs alternating side chains to opposite faces of the cycle.

The search space includes 6-mer, 8-mer, and 9-mer scaffolds (peptoid data bank codes 07AA1-6-C, 07AA2-8-C[26], and 12AC2-9-C[27] respectively) as the backbone and 3-aminopropyl-1-thiol groups as side chains of residues (Fig. 3). Low energy matches were commonly found to be comprised of alternating residue positions, or sequential positions on the narrow end of the macrocycle.

We sampled 8 dihedral angles per atom with different lengths of side chains(n = number of carbon atoms), different error values. We recorded the run time to find the first valid target polygon on Intel Core 3.5 GHz. Results recorded in seconds of CPU run time.

### 3.3 Long loop closure

Loop closure is an increasingly researched field in protein design mainly due to impactful applications including antibody designs [9, 10]. The abstraction of the problem can be described as follows. Given two fixed points in 3D called pivots and two vectors (the take-off vectors), construct the loop from pivot 1 to pivot 2 with k residues with the type N-Cα-C such that the loop has a low energy and it fits in the designated space. Typically the number of residues k ranges from 9 to 17. The difficulties of the problem using a direct computation stem from exponential growth in the number of possible loop conformations as a function of loop length. As illustrated in Fig. 5.5, we divide the loop into two semi-loops by the midpoint or the closest point to the midpoint between two residues. The designated space where the loop resides within can be discretised into cubes of a certain size (the “lattice space” in Fig. 5.5). We precompute all conformations of a single residue and store the resulting angles and x,y,z coordinates after discretisation and encoded as a unique integer. Then we compute and store the table where two residue conformations can connect appropriately, that is, the end atom of one residue and the beginning atom of the other residue lie in the same cube and the two bonds form an angle within the error bound from the optimal bond angle. Now using the precomputed residue conformations and matching table, we develop the two semi-loops. Let the number of residue conformations be Mr and the number of cubes in the lattice space Mc. After developing each residue, we collapse the end positions that fall into the same cube and sharing the same last bond angle, and store all intermediate results for the purpose of producing final results in backtracking. After the two semi-loops are developed, we have the end atoms of both sides and their spatial intersections. The angles are checked to eliminate from the intersection cubes those that deviate outside the error bound from the optimal bond angle there (Fig. 5.6). Starting from the matched cubes in the middle of the loop, now we backtrack in both sides to the pivots and produce as many results as desired (effectively allowing for efficient sampling of a large number of constraint-compliant loop designs). In the first experiment we computed a 12-residue loop, developing 1000 conformations for each residue and 121 by 121 by 121 cubes in the designated space, setting cube length to 0.1 and maximum bond angle errors to within 0.2 rad. On a 1.3 GHz Intel Core M with 8 GB memory, our algorithm ran a total of 3.6 minutes to produce the first result (see Fig. 5.7 for sample results.). The development of each semi-loop took 82 seconds and the matching in the middle took 20 seconds. Keeping the number of conformations per residue, error bounds and the cube size, we enlarge the number of cubes to 171 by 171 by 171 to compute for 17-residue loops. On a 2.0 GHz Intel Xeon E5-2620 CPU with 128 GB memory, our algorithm ran a total of 35.5 minutes to produce the first result (see Fig. 5.8 for sample results). The development of each semi-loop took 11 minutes and the matching in the middle took 28 seconds.

**Table 1:** Run times for matching different geometric representations of metal binding sites to a library of peptoid (peptidomimetic) scaffolds. Run times are shown in seconds for runs computed on an Intel Core 5 3.5 GHz processor. Run times are shown for three classes of binding site pattern and for various user defined settings (corresponding to different allowable error and approximation ranges in atomic units).

| err | 0.05 |  |  | 0.1 |  |  | 0.5 |  |  |
| --- | --- | --- | --- | --- | --- | --- | --- | --- | --- |
| $\eta$ | 0.1 | 0.2 | 0.3 | 0.1 | 0.2 | 0.3 | 0.1 | 0.2 | 0.3 |
| General Triangle: |  |  |  |  |  |  |  |  |  |
| $n = 2$ | 2.69 | 2.42 | 2.49 | 2.92 | 2.40 | 2.50 | 7.81 | 6.93 | 6.97 |
| $n = 3$ | 76.64 | 73.56 | 67.87 | 106.96 | 101.12 | 91.04 | 1518.47 | 1357.48 | 1178.47 |
| Equilateral triangle: |  |  |  |  |  |  |  |  |  |
| $n = 2$ | 5.43 | 5.12 | 4.78 | 5.41 | 4.47 | 4.37 | 5.29 | 4.10 | 3.93 |
| $n = 3$ | 181.04 | 161.34 | 139.75 | 175.37 | 151.46 | 124.03 | 272.62 | 223.75 | 176.27 |
| General 4-gon: |  |  |  |  |  |  |  |  |  |
| $n = 2$ | 7.63 | 7.06 | 7.15 | 7.75 | 7.33 | 7.47 | 9.67 | 8.67 | 8.82 |
| $n = 3$ | 224.16 | 218.69 | 209.19 | 271.98 | 254.81 | 235.12 | 3262.67 | 2780.21 | 2079.59 |

## 4 Discussion

We have presented an adaptive method for finding matches between target geometric patterns (that represent protein and peptidomimetic design goals) and scaffolds (which can serve as the biosynthetic or organic synthesis method for positioning side chains in the desired/target geometry). In the protein, enzyme, and peptidomimeic design communities, these geometric search tasks are increasingly becoming limiting steps in design processes. This trend will increase as we scale to larger target patterns and as we compare to growing databases of proteins, peptidomimetic structures and other scaffolds. We have tested our adaptive octree method in three realistic design settings (each one adapted from a recent design paper using geometric target-scaffold or geometric matching) and in each case we were able to speed up the required calculation by 100 to 10,000 fold over previous methods. These speedups allow us to replace poorly scaling heuristics with our algorithm and thus guarantee scaling and run times in a wide variety of design tasks. In addition, our algorithm allows for an explicit specification of allowable error rates and mismatches (built into both the search and the initial construction of the core octree data-structure). Future work could include providing a better interface to the specification of error and allowable mismatches, resulting in a mismatch tolerant geometric search (akin to gaps in sequence alignments). Another area for future work would be to adapt our geometric search to a multiple-alignment setting, allowing us, for example, to seed a search and subsequently update the parameters of the search to reflect families of discovered sites on proteins. This would provide an algorithmic framework for iterative construction of functional sites on proteins that would be informed (in a data-driven manner) by geometric variation across discovered functional sites.

An immediate advantage of our improvement in computational efficiency is that it expands, by improving scaling, the range and types of peptidomietic and protein scaffolds that can be explored. For example, our method dramatically increases the maximum pattern (active site to match to potential scaffolds) that can be engrafted via matching. This is important for enzyme design and catalysis design, as full sites that include substrate binding and catalytic sites can include large numbers of side chains (large numbers of component vectors in the template/starting geometric pattern to be matched/searched)[5, 6]. The design of protein binding sites can also involve large target patterns that challenge previous methods. Our work here opens the door to a more efficient approach to designing these larger patterns and also offers better algorithmic guarantees than previous heuristics. Our examples here show (presented above and as supplemental code) integration with the Rosetta design framework and thus demonstrate how one might integrate our method with a very wide variety of design tasks including protein interface antogonist design, protein interface engraftment, enzyme design, peptidomimetic design, and the engraftment of complex metal binding sites onto target proteins[21, 28]. The computational efficiency of our algorithm also enables new approaches where geometric matching is integrated more tightly with design protocols (for example, integrated into inner search loops instead of simply being performed to set up initial poses or discover starting scaffolds for a design run). The code is freely available as a set of python scripts (https://github.com/JiangTian/adaptive-geometric-search-for-protein-design).

## 5 Acknowledgments

The authors thank the NYU high performance IT computing team. We thank Ian Fisk and Nicholas Carriero of the Simons Foundation for discussion and help with computing. The authors acknowledge the support of the Simons Foundation, the NIH, the NSF and NYU for supporting this research, particularly NSF: MCB-1158273, IOS-1339362, MCB-1412232, MCB-1355462, IOS-0922738, MCB-0929338, and NIH: 2R01GM032877-25A1. We also thank Professor Saurabh Ray of NYU Abu Dhabi for his insights into the state of the art in computational geometry.

## A Appendix

**Fig. 4:**
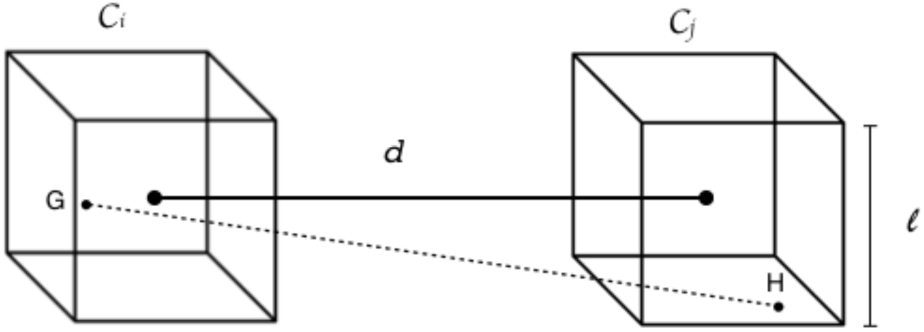
Cubes 𝒞_i_, 𝒞_j_ of size *ℓ* that are *d* distance apart.

### Theorem 1

If 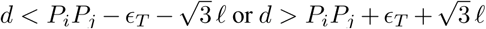, then there are no pairs of points (*G*, *H*) ∈ 𝒞_*i*_ × 𝒞_*j*_ such that |*GH* − *P_i_P_j_*| ≤ *∊_T_*.

*Proof.* For any two points *G* ∈ 𝒞_*i*_, *H* ∈ 𝒞_*j*_ as shown in Figure 4, if *d* < *P_i_P_j_* − *∊_T_* − 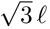, by the triangle inequality we have,

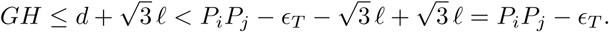

If 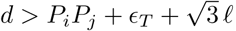, again by the triangle inequality,

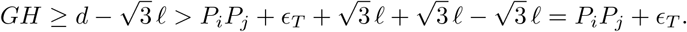

### Theorem 2

If 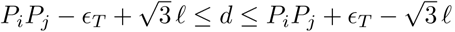, then all pairs of points (*G*, *H*) ∈ 𝒞_*i*_ × 𝒞_*j*_ satisfy |*GH* − *P_i_P_j_*| ≤ *∊_T_*.

*Proof.* As shown in Figure 4, for any points *G* ∈ 𝒞_*i*_, *H* ∈ 𝒞_*j*_, we have 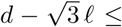 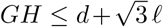. If 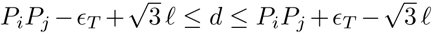 Substituting the tighter bound of *d* on each side of the inequality we have *P_i_P_j_* − *∊_T_* ≤ *GH* ≤ *P_i_P_j_* + *∊_T_*.

### Theorem 3

If we set 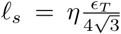 for any 0 < *η* < 1, then the adaptive geometric search algorithm 1 returns all the pairs of points whose distances are within the set [*ℓ*\* − (1 − *η*) *∊_T_*, *ℓ*\* + (1 − *η*)*∊_T_*], and some but possibly not all the pairs of points whose distances are within the set [*ℓ*\* − *∊_T_*, *ℓ*\* − (1 − *η*)*∊_T_*) ∪ (*ℓ*\* + (1 − *η*)*∊_T_*, *ℓ*\* + *∊_T_*].

*Proof.* Let *ℓ_T_* be the length of the leaf cubes. By the definition of *l_s_*, we have *ℓ_T_* < 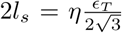. Thus 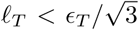 and the sufficient condition A can be tested. If the sufficient condition A is rejected on a pair of cubes 𝒞_1_, 𝒞_2_, then the distance *d* between them satisfies 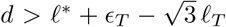 or 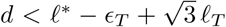. Let *G*, *H* be any two points such that *G* ∈ 𝒞_1_, *H* ∈ 𝒞_2_. By the triangle inequality, we have

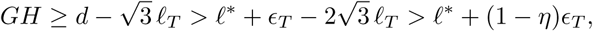

or

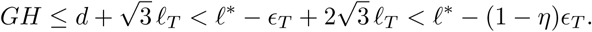

Therefore, in rejecting all pairs of points in 𝒞_1_ × 𝒞_2_ we may have rejected some pairs of points whose distances are within the set [*ℓ*\* − *∊_T_*, *ℓ*\* − (1 − *η*)*∊_T_*) ∪ (*ℓ*\* + (1 − *η*)*∊_T_*, *ℓ*\* + *∊_T_*].

### Lemma 4

Set 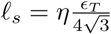 for some 0 < *η* < 1. Then for any cube 𝒞_1_ in an octree *t*_1_, there are at most 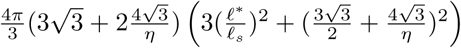 cubes 𝒞_2_ on the same level from another octree *t*_2_ such that (𝒞_1_, 𝒞_2_) are possible pairs, that is, they satisfy the necessary condition 1.

*Proof.* For any cube 𝒞_l_ in *t*_1_, let *∓* be the length of 𝒞_1_. For any possible cube 𝒞_2_ on the same level from *t*_2_, by the necessary condition A the distance between them *d* must satisfy that 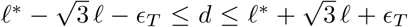. Thus all possible cubes 𝒞_2_ must be contained in the spherical shell 𝒮 of inner radius 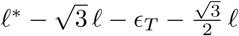 and outer radius 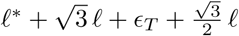 (see Figure 5). Since there are no overlapping cubes on the same level in *t*_2_, the maximum number of the possible cubes *n_m_* satisfies

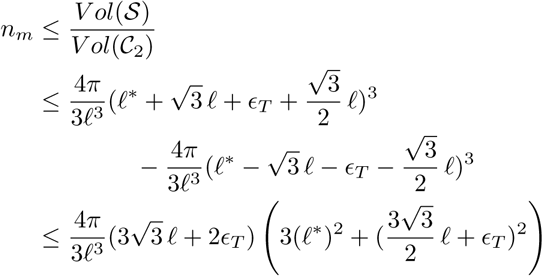

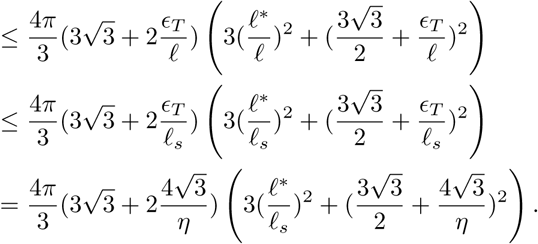

### Theorem 5

Recall that *ℓ*_0_ denotes the initial cube length and the minimum cube length 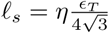. Let *n_m_* be defined as in Lemma A. Then the time complexity of the adaptive search part of Algorithm 1 is 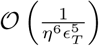.

*Proof.* Let *d* be the depth of the octrees *t*_1_, *t*_2_. Let *ψ_k_*(*t*) be the number of nodes on the *k*-th level in the octree *t*_1_. Recall that *ℓ*_0_ denotes the length of the root cubes of the octrees *t*_1_, *t*_2_. Since all the cubes have the minimum length *ℓ_s_*, we have *ℓ*_0_/2^*d*^ ≥ *ℓ_s_*, or *d* ≤ log_2_ (*ℓ*_0_/*ℓ_s_*). By Lemma 4, the total number of computations 𝒩 satisfies

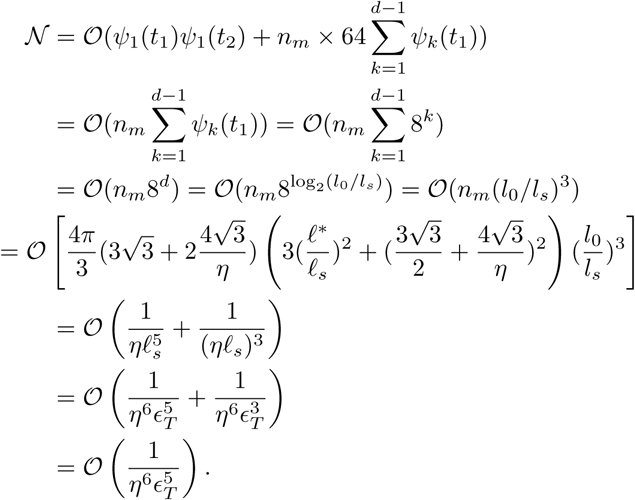

**Fig. 5:**
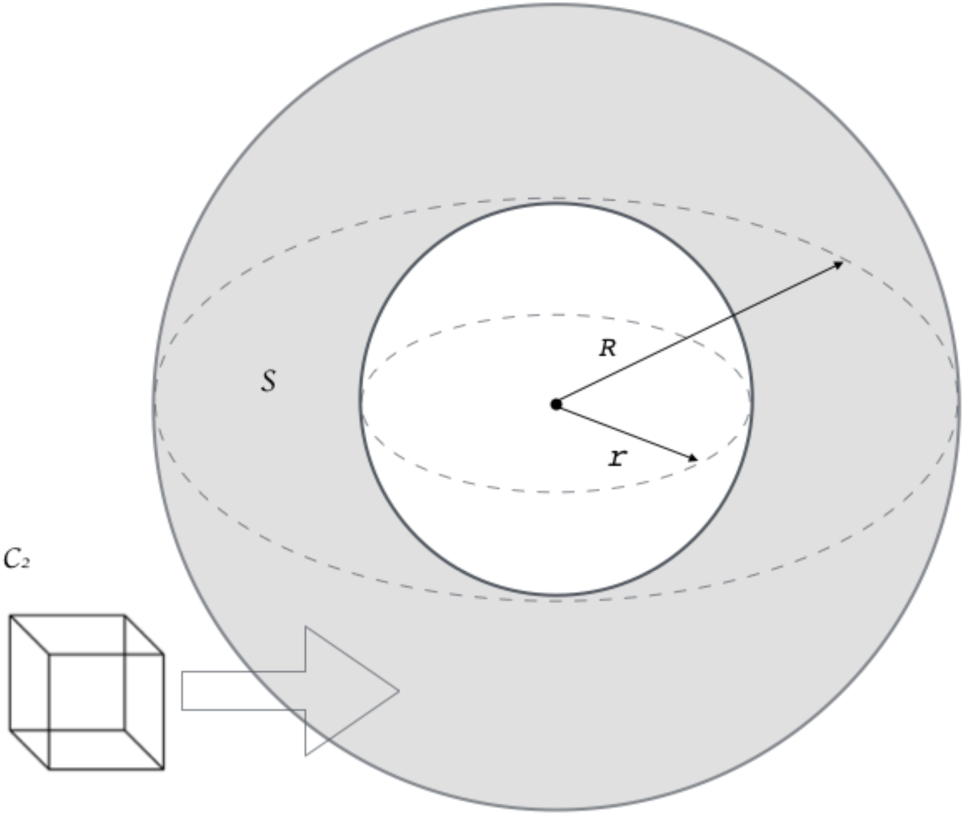
An illustration for Lemma A. The outer radius 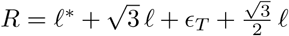, and the inner radius 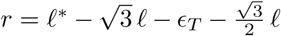. Let 𝓢 denote the spherical shell (in shade). How many cubes 𝒞_2_ can fit into 𝓢?

